# Structural Tuning of HEWL Amyloid Polymorphs Enhances Antibacterial Activity Against Gram-Positive and Gram-Negative Pathogens

**DOI:** 10.64898/2026.07.13.738270

**Authors:** Sanjay K. Metkar, Annie Scutte, Jamel Ali, Ayyalusamy Ramamoorthy

## Abstract

Amyloid fibrils are traditionally associated with protein misfolding disorders; however, increasing evidence indicates that they can also perform beneficial biological functions, including antimicrobial defense. Here, we investigated whether structurally distinct amyloid polymorphs of hen egg white lysozyme (HEWL) exhibit enhanced antibacterial activity compared with the native protein. HEWL was converted into two amyloid polymorphs, flexible fibrils (FFs) and rigid fibrils (RFs), and their antibacterial activities were evaluated against the Gram-positive bacterium *Staphylococcus aureus* and the Gram-negative bacteria *Escherichia coli* (Top10) and *Salmonella Typhimurium*. Fibril formation was confirmed by circular dichroism (CD) spectroscopy, thioflavin T (ThT) fluorescence, and transmission electron microscopy (TEM), demonstrating morphologically distinct amyloid assemblies with different secondary-structure organizations. Fluorescence-based bacterial growth assays showed that native HEWL exhibited only moderate antibacterial activity, whereas both amyloid polymorphs produced potent, concentration-dependent bacterial growth inhibition. FFs and RFs consistently displayed greater antibacterial efficacy than native HEWL across all tested strains, with FFs exhibiting slightly stronger activity against *S. Typhimurium*. At concentrations of 600–800 μM, FFs achieved >90% growth inhibition for all bacterial species examined. Cytotoxicity studies using SH-SY5Y human neuroblastoma cells demonstrated minimal toxicity for native HEWL, modest effects for FFs, and substantially greater toxicity for RFs, indicating that amyloid polymorphism influences both antimicrobial activity and mammalian cell compatibility. Collectively, these findings establish a direct relationship between amyloid structure, antibacterial efficacy, and cytotoxicity. The combination of potent antibacterial activity and relatively low cytotoxicity identifies FFs as a promising functional amyloid biomaterial for the development of next-generation antimicrobial materials.

## 1. Introduction

The overuse of antibiotics has contributed to the emergence of antimicrobial resistance (AMR) in a wide range of microorganisms, including bacteria, viruses, and fungi. AMR poses a serious global health challenge, affecting healthcare systems in both low-income and developed countries. Without effective intervention, AMR is projected to cause up to 10 million deaths annually by 2050 [1]. The development of new antibiotics remains limited because high production costs and complex regulatory approval processes reduce their attractiveness to pharmaceutical companies [2]. Consequently, there is an urgent need for alternative strategies to combat AMR. Broad-spectrum antimicrobial materials, such as antimicrobial peptides, are particularly promising because they can act against diverse range of microorganisms.

The antimicrobial activity of lysozyme in hen egg white (HEWL) and human nasal secretions was first reported in the early 20^th^ century and arises from its ability to cleave the peptidoglycan layer of Gram-positive bacterial cell walls, leading to bacterial lysis and death [3]. Subsequent studies have shown that mutated and heat-denatured lysozyme retain antimicrobial activity even in the absence of catalytic active site [4]. The non-catalytic activity is attributed to structural motifs rich in positively charged amino acid residues that interact with negatively charged bacterial cell walls or membranes, resulting in membrane disruption and cell lysis [5, 6]. In addition, hydrophobic residues may further enhance interactions with microbial membranes [7].

Lysozyme is known to form amyloid aggregates, which are self-assembled fibrils characterized by β-sheet-rich structures stabilized by hydrogen bonding [8]. The ability to generate these amyloids from inexpensive commercial protein sources such as HEWL makes them promising bio-derived nanomaterials with a wide range of potential applications [9]. However, lysozyme-based biomaterials often lose bioactivity during fabrication because acidification and high-temperature conditions cause lysozyme to unfold and hydrolyze, producing rigid fibrils (RFs) that lack enzymatic activity [10]. Interestingly, human lysozyme and HEWL, which share ∼76% sequence identity, can form two distinct amyloid fibril types: one flexible and thermally reversible, and the other rigid and irreversible, without any change in primary sequence [11]. Unlike conventional rigid fibrils, flexible fibrils form through rapid heat shock at neutral pH, a process that induces partial unfolding without hydrolysis and preserved lysozyme functional domains. These fibrils contain a short β-sheet-rich core that loosely confines partially unfolded lysozyme molecules, resulting in significantly lower rigidity than that of conventional rigid fibrils. Notably, flexible fibrils (FFs) exhibit thermal reversibility, dissociating into enzymatically active lysozyme monomers at approximately 50°C and reassembling into fibrils upon reheating [10]. In this study, we hypothesized that FFs, which preserve lysozyme functional domains while containing a β-sheet–rich amyloid core, could exhibit enhanced antibacterial activity than native lysozyme and rigid amyloid fibrils of HEWL. To evaluate the antibacterial potential of HEWL amyloid polymorphs, we selected three clinically relevant bacterial pathogens: *Staphylococcus aureus*, *Escherichia coli*, and *Salmonella* Typhimurium. *S. aureus* is a major Gram-positive pathogen associated with skin and soft tissue infections, pneumonia, and bloodstream infections [12]. *E. coli* (Top 10) and *S.* Typhimurium are Gram-negative bacteria commonly implicated in urinary tract infections, gastrointestinal diseases, foodborne illnesses, and systemic infections [13, 14]. The increasing prevalence of antibiotic-resistant strains among these pathogens highlights the urgent need for alternative antimicrobial strategies. Accordingly, this study characterizes HEWL aggregation kinetics by ThT fluorescence, fibril morphology by TEM, secondary structure by CD spectroscopy, antibacterial activity against clinically relevant bacterial pathogens, and cytotoxicity toward mammalian cells to determine the therapeutic potential of distinct HEWL amyloid polymorphs.

## 2. Material and Methods

### 2.1 Materials

Sodium chloride (NaCl) and sodium phosphate were purchased from Fisher Scientific (Hampton, NH, USA). Copper grids used for transmission electron microscopy were supplied by Millipore Sigma (Burlington, MA, USA), and Uranyless Stain was purchased from Electron Microscopy Sciences. Thioflavin T (ThT) fluorescence dye was obtained from Millipore Sigma (Burlington, MA, USA). Chicken egg white lysozyme (Catalog No. J60701.14) was purchased from Thermo Scientific Chemicals (USA) and used as received. SH-SY5Y human neuroblastoma cells were obtained from the American Type Culture Collection (ATCC, Manassas, VA, USA). Dulbecco’s Modified Eagle Medium (DMEM), fetal bovine serum (FBS), penicillin–streptomycin, phosphate-buffered saline (PBS), and 0.25% trypsin–EDTA were purchased from Corning Inc. (Corning, NY, USA). Cell viability was assessed using the CyQUANT™ XTT Cell Viability Assay (Cat. No. X12223; Invitrogen™, Thermo Fisher Scientific). Fluorescence imaging was performed using BioTracker™ 555 Orange Cytoplasmic Membrane Dye (SCT107; Sigma-Aldrich, St. Louis, MO, USA), 4% paraformaldehyde in PBS (Cat. No. J61899.AK; Thermo Fisher Scientific), and Fluoromount-G™ Mounting Medium containing DAPI (Invitrogen™, Thermo Fisher Scientific). 8-well chamber slides (CELLTREAT Scientific Products, Pepperell, MA, USA; Cat. No. 50-114-9054), glass coverslips (VWR®, Avantor, Radnor, PA, USA), and sterile flat-bottom 96-well tissue culture plates (Fisherbrand™, Fisher Scientific, Waltham, MA, USA) were used for cell culture experiments.

### 2.2 Preparation of HEWL Amyloid Fibrils

FFs and RFs HEWL amyloid fibrils were prepared following the protocol described by Frey et al. (2024) with slight modifications. FFs were prepared by incubating a solution containing 20 mg/ml lysozyme in sodium phosphate buffer (NaPi, pH 7.0), 20 mM TCEP, and 10 mM NaCl at 85 °C for 5 minutes with continuous shaking at 300 rpm. RFs were prepared with the same composition as FFs but incubated at 85 °C for 3 hours with shaking at 300 rpm. Prior to fibril formation, excess tris(2-carboxyethyl) phosphine (TCEP) was removed from the reduced lysozyme solution by centrifugation using an Amicon Ultra-0.5 centrifugal filter unit (10 kDa molecular weight cut-off, UFC501008, MilliporeSigma, Burlington, MA, USA), and the retentate containing purified reduced lysozyme was collected for fibril preparation.

### 2.3 Morphological Characterization of HEWL fibrils by TEM

The morphology of FFs and RFs of HEWL was examined by TEM. Freshly prepared FFs and RFs (10 µL) were adsorbed onto 300-mesh Formvar/carbon-supported copper grids (catalog #TEM-FCF300CU, MilliporeSigma, Burlington, MA, USA) and allowed to adsorb for 10 minutes at room temperature. Excess sample was removed by blotting, and grids were negatively stained with 10 µL of Uranyless stain (catalog #22409, Electron Microscopy Sciences, Hatfield, PA, USA) for 1 minute. Excess stain was carefully removed using blotting paper, and grids were air-dried at room temperature for 10 minutes prior to imaging. All FFs and RFs were imaged using a Hitachi High-Tech HT7800 transmission electron microscope (Minato-ku, Tokyo, Japan) at an acceleration voltage of 100 kV. To ensure representative sampling, images were acquired from a minimum of three independent grid regions for each FFs and RFs preparation.

### 2.4 Thioflavin T (ThT) Fluorescence Assay

ThT fluorescence was used to measure amyloid fibril formation. HEWL samples including monomers, FFs and RFs (100 µM) were mixed with 30 µM ThT in 10 mM NaPi buffer (pH 7.0) and added to black 384-well plates. Plates were incubated at 37 °C with shaking at 700 rpm. Fluorescence was measured using a Varioskan ALF reader at 440 nm excitation and 480 nm emission. Buffer-only and ThT-only wells were used as controls. Data were background-corrected and averaged from at least three independent experiments.

### 2.5 Conformational Analysis of HEWL Fibril Polymorphs by Far-UV CD

Secondary structure of FFs and RFs of HEWL was analyzed by CD spectroscopy using a Chirascan spectropolarimeter (Applied Photophysics, Leatherhead, UK). Samples were diluted to 30 µM in 10 mM NaPi (pH 7.0), and spectra were recorded over the far-UV range (200–250 nm) at room temperature. Three scans were averaged per sample following baseline subtraction, and spectra were visualized using GraphPad Prism (v11.0.2, GraphPad Software, Boston, MA, USA).

### 2.6 Antimicrobial Activity of FFs, RFs, and Native HEWL

#### 2.6.1 Bacterial Strains and Culture Conditions

To evaluate the broad-spectrum antimicrobial potential of FFs, RFs ana Native HEWL, a panel of representative Gram-positive and Gram-negative bacteria were employed: *Escherichia coli* TOP10 (K12, Gram-negative), *Salmonella* Typhimurium (CDC 6516-60, Gram-negative), and *Staphylococcus aureus* (155554A, Gram-positive). All strains were cultured in Luria–Bertani (LB) broth (1% tryptone, 0.5% yeast extract, 1% NaCl, pH 7.4) at 37 °C under aerobic conditions with orbital shaking at 150 rpm. BSL-2 organisms were handled in a certified Class II biosafety cabinet in accordance with institutional biosafety guidelines.

#### 2.6.2 Resazurin-Based Bacterial Viability Assay

The antimicrobial activity of FFs and RFs of HEWL was assessed using a resazurin (Alamar Blue) reduction assay, which quantifies the metabolic activity of viable bacteria. Overnight cultures were harvested by centrifugation, washed twice with sterile phosphate-buffered saline (PBS, pH 7.4), and resuspended in PBS to a standardized cell density (OD_₆₀₀_ ≈ 0.1). Bacterial suspensions were dispensed into black, flat-bottom 384-well microplates (Corning) and treated with FFs or RFs at the indicated concentrations. Monomeric HEWL was included as a non-fibrillar control at matched concentrations to distinguish the contribution of the fibrillar state from that of the native protein, while untreated suspensions in PBS served as viability controls. Plates were incubated at 37 °C with orbital shaking at 150 rpm for 12 h. As PBS does not support bacterial proliferation, this format isolates the direct bactericidal effect of the fibrils from growth-dependent variables.

Following exposure, resazurin reagent (Alamar Blue) was added to each well at 10% (v/v), and plates were further incubated at 37 °C in the dark until fluorescence developed. The metabolic reduction of non-fluorescent resazurin to fluorescent resorufin was measured using a microplate reader (λex = 560 nm; λem = 590 nm). PBS-only and fibril-only wells were included as background controls to correct for autofluorescence and non-specific reagent reduction. Fluorescence values were background-subtracted and normalized to untreated controls to determine relative bacterial viability following fibril exposure. Control experiments confirmed that RFs and FFs alone did not produce significant Alamar Blue fluorescence in the absence of bacteria (Supplementary Figure S1), indicating negligible assay interference.

### 2.7. Cell Culture, Cytotoxicity Assessment, and Fluorescence Imaging

#### 2.7.1 Cell Culture

SH-SY5Y human neuroblastoma cells were cultured in Dulbecco’s Modified Eagle Medium (DMEM) supplemented with 10% fetal bovine serum (FBS) and 1% penicillin–streptomycin and maintained at 37 °C in a humidified incubator containing 5% CO_₂_. Cells were routinely passaged upon reaching 70–80% confluency. Briefly, the culture medium was removed, cells were washed with phosphate-buffered saline (PBS), detached using 0.25% trypsin–EDTA, collected by centrifugation, resuspended in fresh complete medium, and seeded into new culture vessels for subsequent experiments.

#### 2.7.2 XTT Cell Viability Assay

For cytotoxicity assessment, SH-SY5Y cells were seeded into 96-well tissue culture plates at a density of 1 × 10 cells per well and allowed to adhere for 24 h. Cells were then treated with native hen HEWL, FFs, or RFs of HEWL at final concentrations ranging from 100 to 800 µM for 72 h. Following treatment, cell viability was determined using the CyQUANT™ XTT Cell Viability Assay (Cat. No. X12223; Invitrogen™) according to the manufacturer’s instructions. XTT reagent was added directly to each well, and absorbance was measured at 450 nm using Varioskan™ ALF Multimode Microplate Reader (Thermo Fisher Scientific, Waltham, MA, USA). Cell viability was calculated relative to untreated control cells using the following equation:

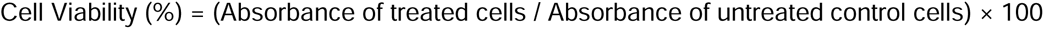

#### 2.7.3 Fluorescence Imaging

For fluorescence microscopy, SH-SY5Y cells were seeded into 8-well chamber slides (CELLTREAT Scientific Products; Cat. No. 50-114-9054) at a density of 1 × 10 cells per well and allowed to adhere for 24 h. Cells were subsequently treated with native HEWL, FFs, or RFs at concentrations ranging from 100 to 800 µM for 48 h. Following treatment, live cells were stained with BioTracker™ 555 Orange Cytoplasmic Membrane Dye (SCT107; Sigma-Aldrich, St. Louis, MO, USA) diluted 1:1000 in complete culture medium and incubated for 30 min at 37 °C, according to the manufacturer’s instructions to visualize plasma membrane morphology. Cells were then washed with PBS and fixed with 4% paraformaldehyde in PBS (Cat. No. J61899.AK; Thermo Fisher Scientific) for 5 min at room temperature. After fixation, cells were washed three times with PBS and mounted using Fluoromount-G™ Mounting Medium containing DAPI (Invitrogen™, Thermo Fisher Scientific, Waltham, MA, USA). Coverslips were carefully placed onto the samples, and the slides were stored overnight at 4 °C before imaging. Fluorescence images were acquired using an Invitrogen™ EVOS™ M7000 Imaging System (Carlsbad, CA, USA) equipped with a 20× objective. Images were collected using the red fluorescence channel to visualize BioTracker™ 555-labeled plasma membranes and the blue fluorescence channel to visualize DAPI-stained nuclei. Representative images were acquired under identical imaging parameters to qualitatively compare treatment-induced changes in plasma membrane morphology and nuclear integrity relative to untreated control cells.

## 3 Results

### 3.1 Structural and Morphological Characterization of Native HEWL and Amyloid Fibril Polymorphs

Figure 1 compares the structural and morphological characteristics of native HEWL with two amyloid forms: FFs and RFs. CD spectroscopy was used to examine changes in secondary structure. Native HEWL shows a characteristic negative band around ∼208–215 nm (Figure 1a), which is typical of proteins with a predominantly α-helical structure, indicating that the protein is properly folded in its native state. In contrast, both FFs and RFs display a shift of the negative minimum toward ∼218–220 nm (Figure 1b-c), which is a hallmark of β-sheet–rich structures commonly observed in amyloid fibrils. The signal for RFs is deeper than that for FFs, suggesting that RFs contain a more ordered and extensive β-sheet arrangement, whereas FFs retain some structural flexibility or less compact β-sheet organization.

**Figure 1.**
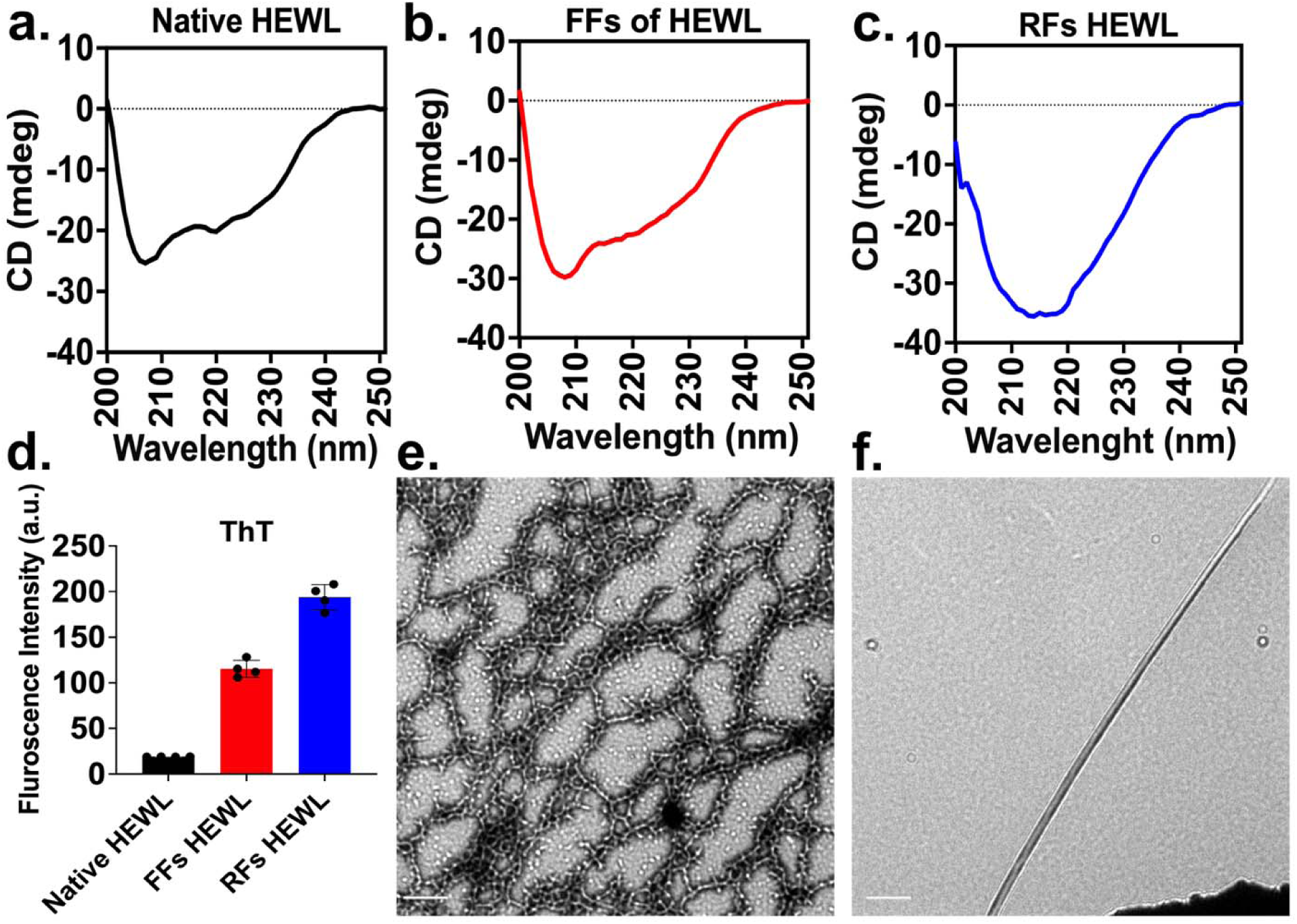
Structural and morphological characterization of lysozyme and its fibrillar forms. (a–c) Far-UV circular dichroism (CD) spectra of (a) native HEWL, (b) FFs, and (c) RFs. Native HEWL shows a spectrum characteristic of an α-helical structure, whereas both FF and RF exhibit a minimum around ∼218 nm, indicating the presence of β-sheet–rich amyloid structures. RF shows a stronger β-sheet signal compared to FF, suggesting a more ordered fibrillar structure. (d) ThT fluorescence assay showing increased fluorescence intensity for FFs and RFs compared to native HEWL, confirming amyloid formation, with RFs displaying the highest fluorescence. (e–f) TEM images of lysozyme fibrils showing (e) FFs with curved, network-like morphology and (f) RFs with long, straight (Scale bar = 200 nm), and well-defined fibrils characteristic of mature amyloid structures.

The formation of amyloid structures was further confirmed using ThT fluorescence assay, which selectively binds to β-sheet–rich amyloid fibrils and increases fluorescence intensity. Native HEWL shows very low fluorescence, indicating the absence of amyloid structures. In comparison, FFs exhibit a moderate increase in fluorescence intensity, suggesting the formation of amyloid-like assemblies with partial β-sheet organization. RFs show the highest fluorescence intensity, indicating a greater degree of β-sheet stacking and more mature amyloid fibril formation (Figure 1d). These results are consistent with the CD data and confirm that RFs possess a higher level of structural order compared to flexible fibrils.

TEM images further reveal distinct morphological differences between the two fibril types. FFs appear as interconnected, curved, and network-like structures, indicating a relatively dynamic and less rigid organization (Figure 1e). These fibrils often show branching and entanglement, suggesting that they consist of loosely packed protofilaments. In contrast, RFs appear as long, straight, and well-defined needle-like structures, characteristic of mature amyloid fibrils with strong intermolecular β-sheet interactions (Figure 1f). Overall, these results demonstrate that while both FFs and RFs adopt amyloid β-sheet structures, FFs represent a less ordered and more dynamic amyloid state, whereas RFs correspond to highly ordered, structurally stable amyloid fibrils.

### 3.2 FFs and RFs of HEWL Polymorphs Exhibit Antibacterial Activity Against Gram positive *Staphylococcus aureus*

The antibacterial activities of native HEWL, FFs, and RFs were evaluated against *Staphylococcus aureus* using a fluorescence-based growth assay (Figure 2). Native HEWL exhibited a concentration-dependent inhibition of bacterial growth, with increasing protein concentration resulting in progressively lower fluorescence intensities relative to the untreated control (Figure 2a). However, bacterial growth remained detectable even at the highest concentration tested (800 µM), indicating only partial growth suppression. In contrast, FFs displayed markedly enhanced antibacterial activity compared with native HEWL (Figure 2b). At concentrations of 400–800 µM, FFs induced a rapid and sustained reduction in fluorescence intensity, indicating strong inhibition of bacterial proliferation. Notably, treatment with 600 and 800 µM FFs nearly abolished bacterial growth throughout the incubation period, demonstrating a substantial gain in antibacterial potency following its amyloid conversion.

**Figure 2.**
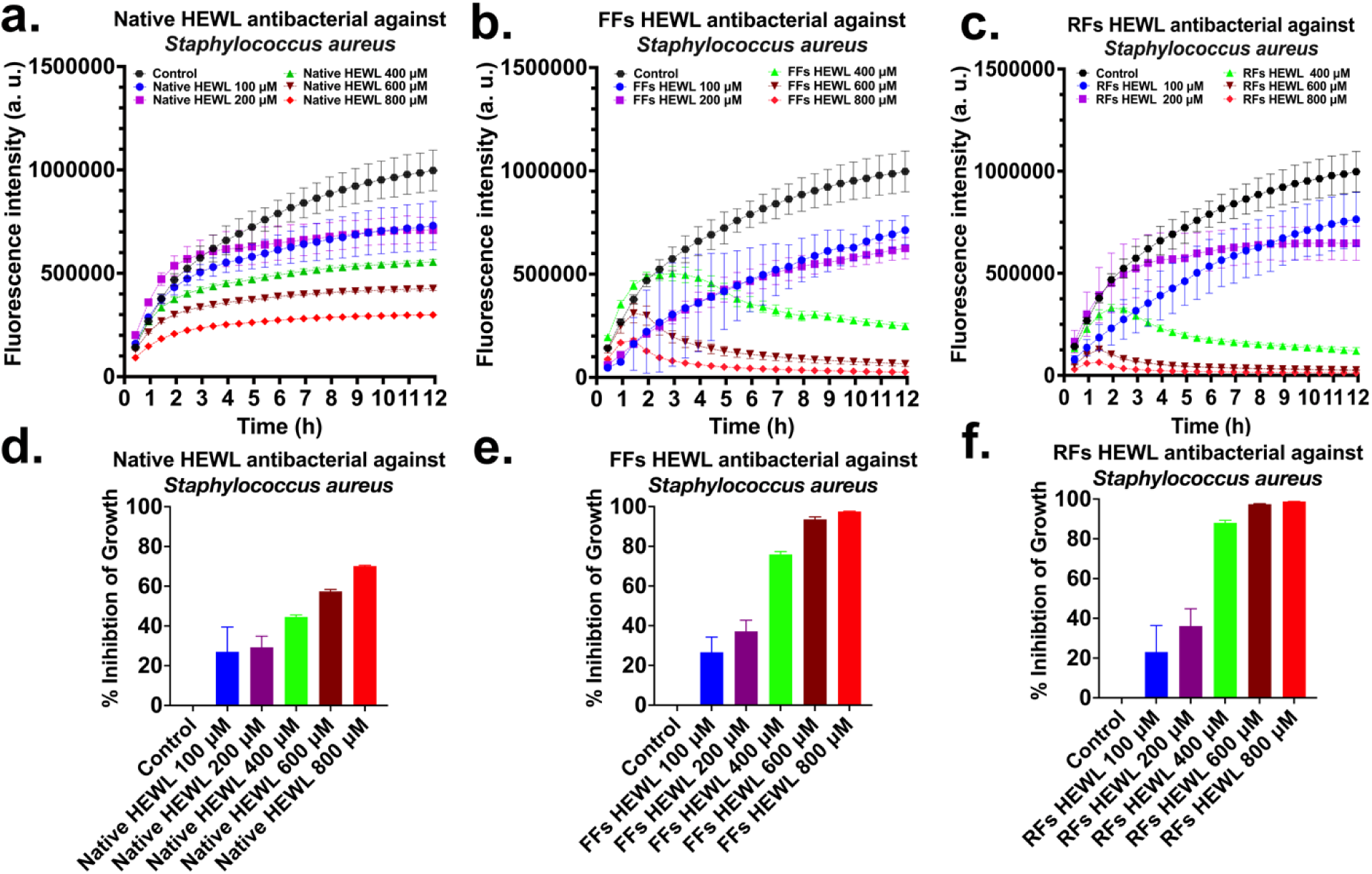
Antibacterial activity of native, FFs, and RFs HEWL against the Gram-positive bacteria *Staphylococcus aureus* measured using the Alamar Blue assay. Bacterial growth kinetics were monitored by fluorescence intensity over 12 h. (a) *Staphylococcus aureus* treated with native HEWL (a), FFs HEWL (b), and RFs HEWL (c). (d–f) Percentage growth inhibition at 12 h relative to untreated controls. FFs and RFs exhibited markedly greater antibacterial activity than native HEWL, with RFs showing the strongest inhibition at higher concentrations. Data are presented as mean ± SD (n = 5). Proteins were tested at 100–800 μM. Control represents untreated bacteria. Reduced fluorescence indicates growth inhibition. Data represent mean ± SD from independent experiments.

The enhanced activity of FFs may arise from their increased surface area, conformational flexibility, and greater capacity to interact with and disrupt bacterial cell envelopes. RFs also inhibited *S. aureus* growth in a concentration-dependent manner (Figure 2c). While RFs exhibited greater antibacterial activity than native HEWL, their efficacy was nearly similar than that observed for FFs at equivalent concentrations. The reduced activity of RFs may reflect their highly ordered and compact fibrillar architecture, which could limit interactions with bacterial membranes relative to the more dynamic FFs structures. Collectively, these findings demonstrate that amyloid polymorphism profoundly influences the antimicrobial properties of HEWL.

To quantitatively compare antibacterial activity, fluorescence intensities measured at ∼12 h endpoint were normalized to the untreated control and expressed as percentage inhibition of bacterial growth (Figure 2 d-f). Native HEWL produced a moderate reduction in bacterial growth across all tested concentrations (Figure 2a). Endpoint inhibition analysis (Figure 2d) showed approximately 27%, 29%, 45%, 57%, and 70% inhibition at 100, 200, 400, 600, and 800 µM, respectively. Although bacterial growth was suppressed relative to the untreated control, substantial proliferation remained evident even at the highest concentration tested. In contrast, FFs displayed markedly enhanced antibacterial activity (Figure 2b). Growth inhibition increased sharply with concentration, particularly above 400 µM. Quantification of endpoint inhibition (Figure 2e) revealed approximately 26%, 37%, 76%, 93%, and 97% inhibition at 100, 200, 400, 600, and 800 µM, respectively. At 600 and 800 µM, FFs nearly abolished bacterial growth throughout the incubation period. RFs exhibited the strongest antibacterial activity among all HEWL formulations (Figure 2b,2e).

Bacterial proliferation was substantially reduced at concentrations ≥400 µM, and almost complete suppression was observed at 600 and 800 µM. Endpoint analysis (Figure 2f) demonstrated approximately 23%, 36%, 88%, 97%, and 99% inhibition at 100, 200, 400, 600, and 800 µM, respectively. Comparison of the inhibition profiles shown in Figures 2d–f revealed a clear enhancement in antibacterial efficacy following amyloid fibril formation. At 400 µM, FFs and RFs achieved approximately 76% and 88% inhibition, respectively, compared with only 45% inhibition for native HEWL. However, the difference between FFs and RFs became progressively smaller at higher concentrations, with both polymorphs producing >93% inhibition at 600 µM and >97% inhibition at 800 µM (Figures 2 e,f). These findings indicate that amyloid assembly significantly enhances the intrinsic antibacterial activity of HEWL, while both fibrillar polymorphs exhibit comparable antibacterial efficacy at higher concentrations.

### 3.3 Morphology-Dependent Antibacterial Activity of HEWL Amyloid Polymorphs Against Gram-Negative Bacteria *Escherichia coli* and *Salmonella typhimurium*

To investigate the impact of amyloid polymorphism on the antibacterial properties of HEWL, the activities of native HEWL and its fibrillar polymorphs (FFs and RFs) were evaluated against the Gram-negative pathogen *Escherichia coli* (Top 10) (Figures 3). Bacterial growth was monitored over 12 h using fluorescence-based assays in the presence of increasing concentrations of HEWL and its amyloid polymorphs (100–800 μM). Untreated control cultures exhibited a continuous increase in fluorescence intensity throughout the incubation period, indicating robust bacterial growth. Native HEWL produced a concentration-dependent reduction in bacterial growth, with the greatest suppression observed at 800 μM (Figure 3a). In contrast, FFs markedly inhibited bacterial proliferation at concentrations ≥400 μM, resulting in substantially lower fluorescence intensities compared with the control and native HEWL groups (Figure 3b). Similarly, RFs exhibited strong antibacterial activity, particularly at 600 and 800 μM, where bacterial growth was significantly suppressed throughout the incubation period (Figure 3c).

**Figure 3.**
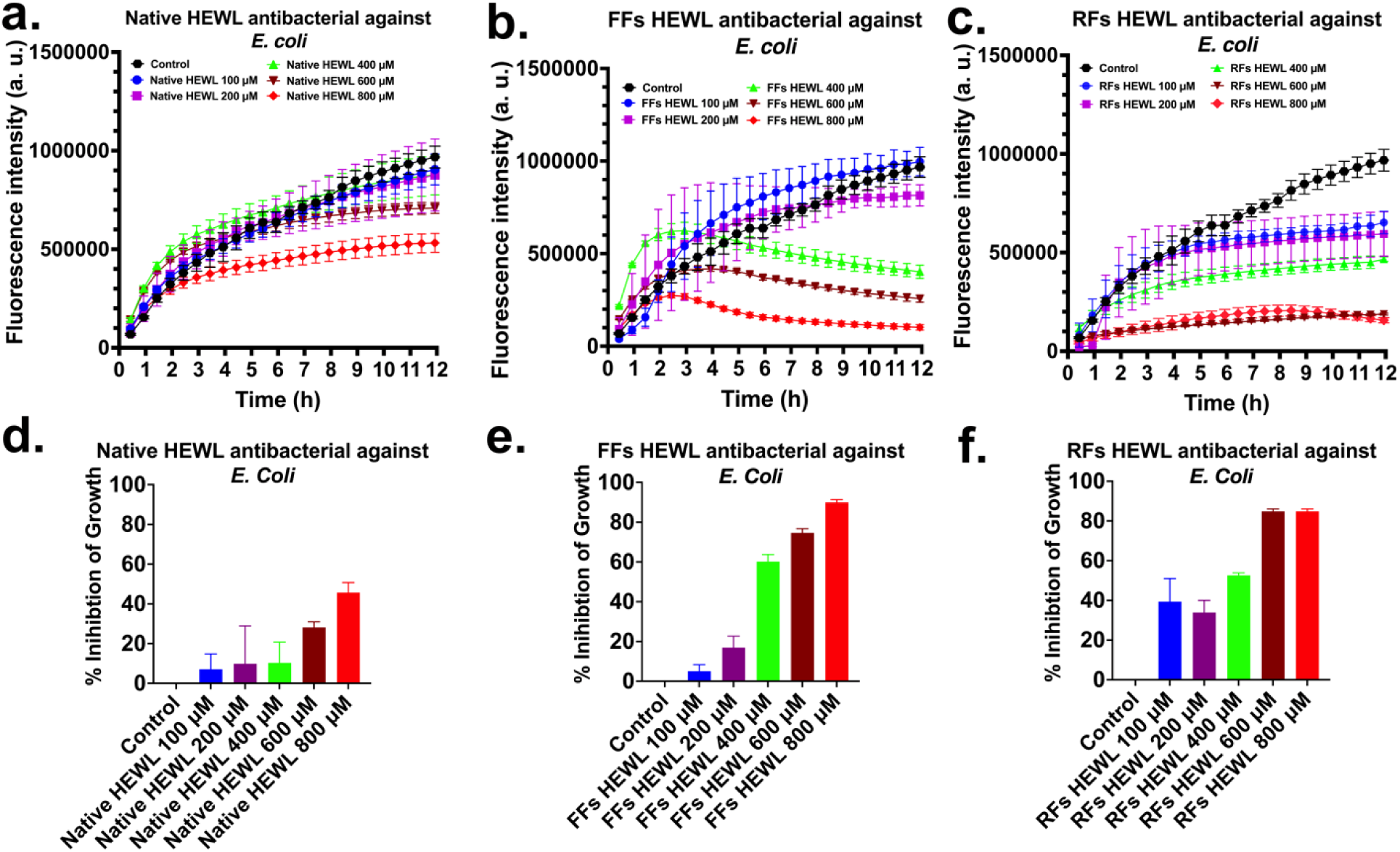
Antibacterial activity of native HEWL and its amyloid polymorphs against *Escherichia coli*. (a–c) Fluorescence-based growth curves of *E. coli* treated with native HEWL, FFs, and RFs, respectively, at concentrations of 100–800 μM over 12 h. (d–f) Percentage inhibition of bacterial growth at the 12 h endpoint relative to untreated controls for native HEWL, FFs, and RFs, respectively. Data are presented as mean ± SD.

To quantitatively compare antibacterial efficacy, fluorescence intensities measured at the 12 h endpoint were normalized to the untreated control and expressed as percentage inhibition of bacterial growth (Figure 3d–f). Native HEWL displayed moderate antibacterial activity, with growth inhibition of approximately 5%, 10%, 10%, 28%, and 45% at 100, 200, 400, 600, and 800 μM, respectively (Figure 3d). FFs exhibited substantially enhanced antibacterial activity compared with native HEWL (Figure 3e). Growth inhibition increased from approximately 6% and 17% at 100 and 200 μM to 60%, 74%, and 90% at 400, 600, and 800 μM, respectively. Similarly, RFs demonstrated potent antibacterial activity against *E. coli* (Figure 3f), achieving approximately 40%, 34%, 53%, 85%, and 86% inhibition at 100, 200, 400, 600, and 800 μM, respectively. Overall, both amyloid polymorphs significantly outperformed native HEWL, particularly at concentrations ≥400 μM. While FFs exhibited the highest inhibition at 800 μM, RFs showed stronger antibacterial activity at lower concentrations. These findings demonstrate that amyloid fibril formation markedly enhances the intrinsic antibacterial properties of HEWL against *E. coli* and that fibril morphology influences antibacterial potency.

The antibacterial activity of native HEWL and its amyloid polymorphs was further evaluated against clinically relevant *Salmonella* Typhimurium using a fluorescence-based growth assay over 12 h (Figure 4a–c). Untreated control cultures showed a continuous increase in fluorescence intensity throughout the incubation period, indicating robust bacterial growth.

**Figure 4.**
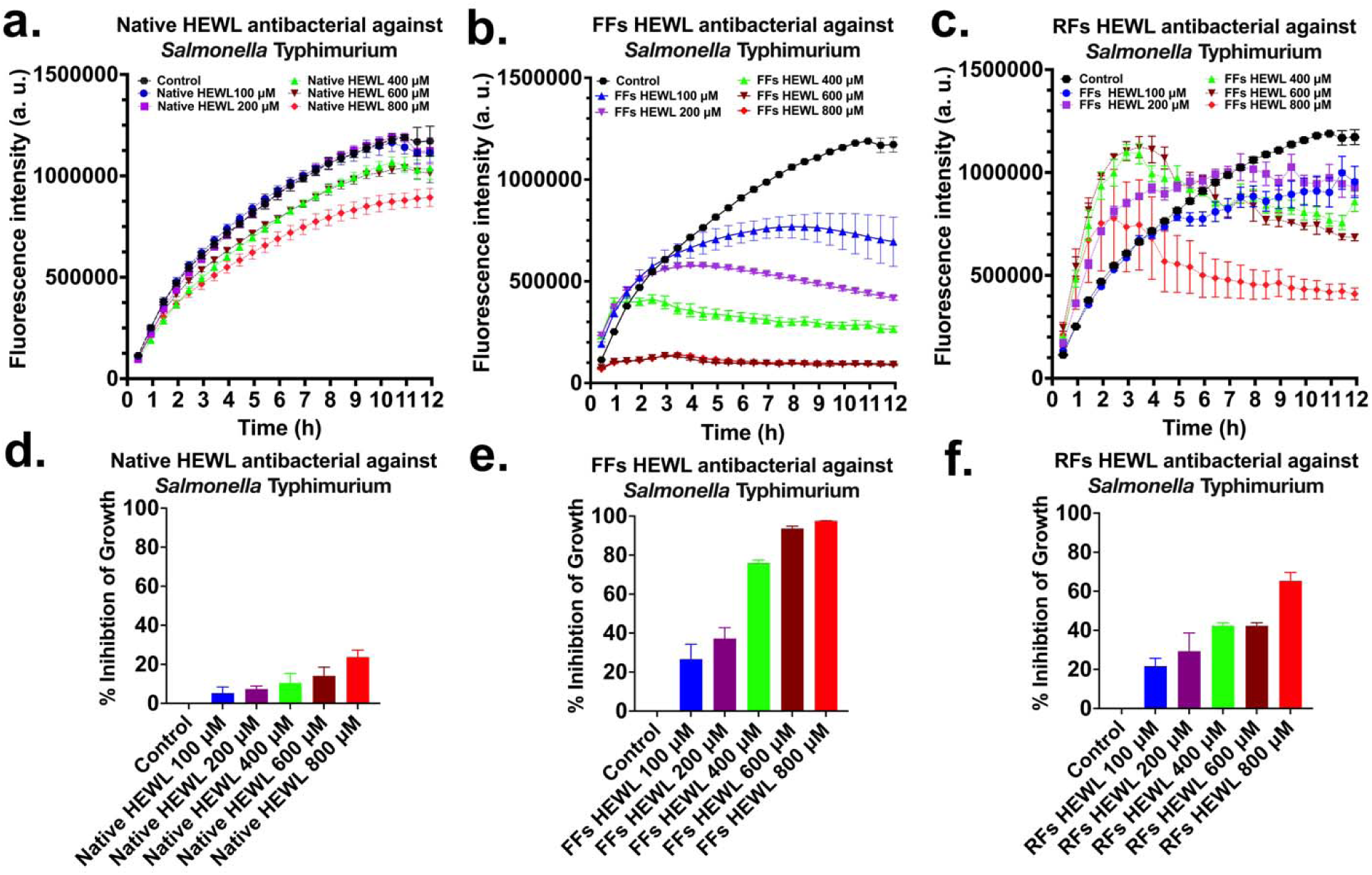
Antibacterial activity of native HEWL and its amyloid polymorphs against *Salmonella* Typhimurium. (a–c) Fluorescence-based growth curves of *Salmonella* Typhimurium treated with native HEWL, FFs, and RFs, respectively, at concentrations of 100–800 μM over 12 h. (d–f) Percentage inhibition of bacterial growth at the 12 h endpoint relative to untreated controls for native HEWL, FFs, and RFs, respectively. Data are presented as mean ± SD.

Native HEWL produced only modest bacterial growth inhibition, with fluorescence intensities remaining comparable to the control at lower concentrations (Figure 4a). In contrast, both FFs and RFs exhibited enhanced antibacterial activity, particularly at concentrations ≥400 μM, resulting in markedly reduced bacterial growth over time (Figure 4b-c). To quantitatively compare antibacterial efficacy, fluorescence intensities measured at the 12 h endpoint were normalized to untreated controls and expressed as percentage inhibition of bacterial growth (Figure 4d–f). Native HEWL displayed relatively weak antibacterial activity, achieving approximately 5%, 6%, 10%, 12%, and 22% inhibition at 100, 200, 400, 600, and 800 μM, respectively (Figure 4d). FFs exhibited substantially enhanced antibacterial activity compared with native HEWL (Figure 4e). Growth inhibition increased from approximately 25% and 37% at 100 and 200 μM to 76%, 95%, and 98% at 400, 600, and 800 μM, respectively. Similarly, RFs demonstrated concentration-dependent antibacterial effects, producing approximately 18%, 23%, 40%, 38%, and 65% inhibition at 100, 200, 400, 600, and 800 μM, respectively (Figure 4f).

Overall, FFs exhibited the strongest antibacterial activity against *S. Typhimurium*, achieving nearly complete growth inhibition at concentrations of 600–800 μM. Although RFs also inhibited bacterial growth more effectively than native HEWL, their antibacterial activity was lower than that of FFs at higher concentrations. These findings demonstrate that amyloid fibril formation significantly enhances the antibacterial properties of HEWL against *S.* Typhimurium, with flexible fibrils displaying the greatest efficacy.

### 3.4 Differential Cytotoxicity of Native HEWL and Amyloid Polymorphs in SH-SY5Y Cells

The cytotoxic effects of native HEWL, FFs, and RFs were evaluated in SH-SY5Y human neuroblastoma cells following 48 h of treatment using the XTT cell viability assay (Figure 5). Native HEWL exhibited minimal cytotoxicity across the concentration range tested (100–800 µM), with cell viability remaining comparable to that of untreated control cells (Figure 5a). Similarly, FFs produced only a modest reduction in cell viability, with slight concentration-dependent decreases observed at higher concentrations while overall viability remained relatively high (Figure 5b). In contrast, RFs displayed substantially greater cytotoxicity than both native HEWL and FFs (Figure 5c). Treatment with RFs resulted in a marked reduction in cell viability at all concentrations tested, with viability decreasing to approximately 40–60% of control levels. These findings indicate that the rigid fibrillar polymorph possesses significantly greater neurotoxic potential than the flexible fibrillar polymorph or the native protein.

**Figure 5.**
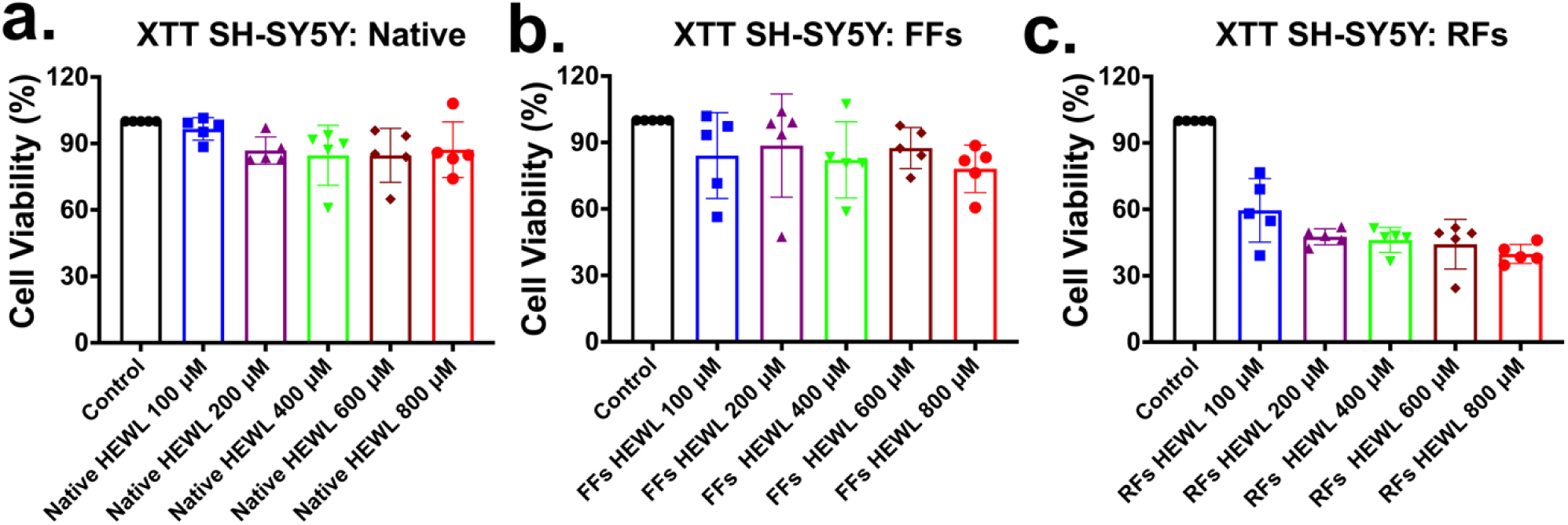
Cytotoxicity of native HEWL and its amyloid polymorphs toward human SH-SY5Y neuroblastoma cells assessed by XTT assay. Cell viability (%) was measured following 48 h treatment with (a) native HEWL, (b) FFs, and (c) RFs at 100–800 μM. Native HEWL did not significantly reduce cell viability. FFs produced mild, concentration-dependent decreases at higher concentrations, while RFs induced a more pronounced and consistent reduction across the full range. Data = mean ± SD (n = 5).

To further investigate the morphological effects of HEWL polymorphs, fluorescence microscopy was performed using BioTracker™ 555 Orange Cytoplasmic Membrane Dye to visualize plasma membrane morphology and DAPI to stain cell nuclei (Figure 6). Untreated control cells displayed intact membrane morphology and uniformly distributed DAPI-stained nuclei. Cells treated with native HEWL showed membrane morphology and nuclear staining patterns comparable to those of the control group across all concentrations, consistent with the minimal cytotoxicity observed in the XTT assay (Figure 6a).

**Figure 6.**
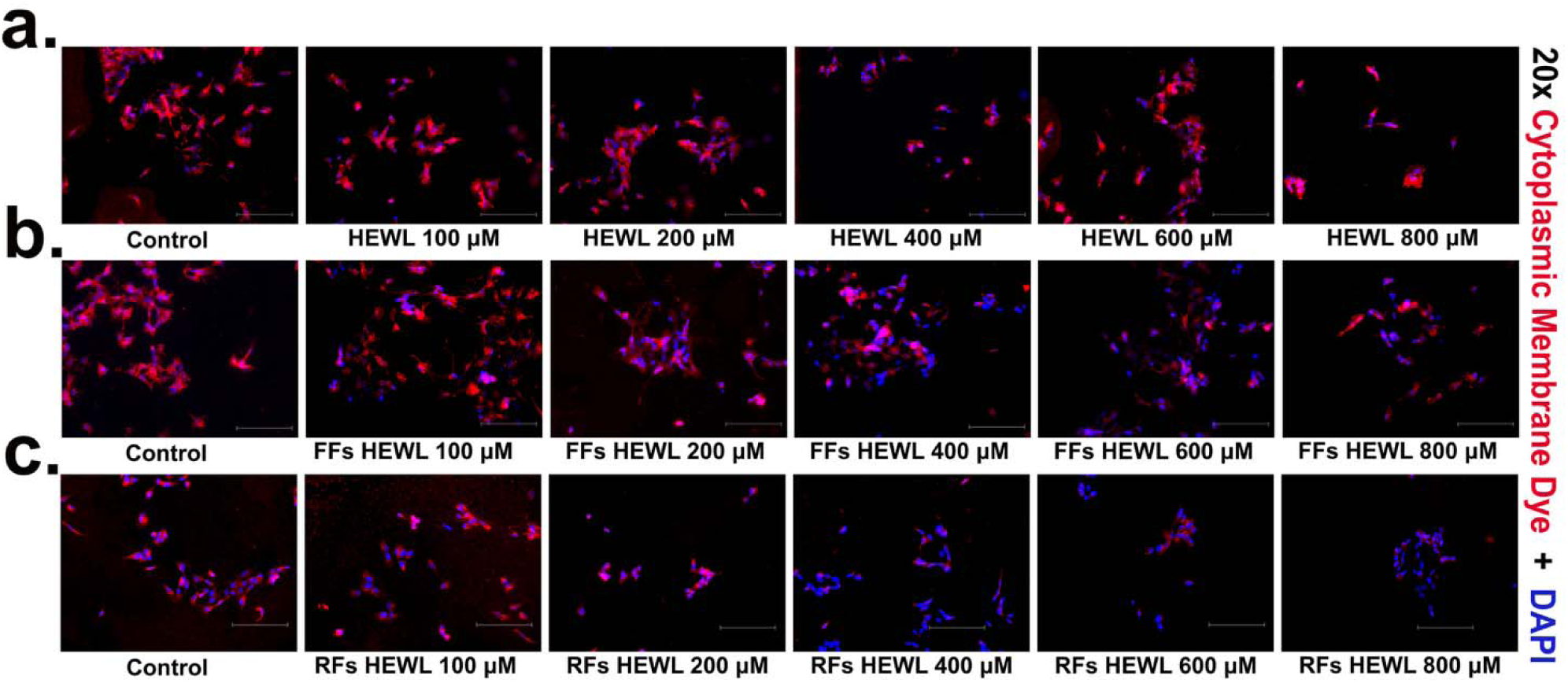
Representative fluorescence microscopy images of SH-SY5Y neuroblastoma cells following treatment with native, FFs, and RFs of HEWL at concentrations ranging from 100 to 800 µM. **(a)** Native HEWL-treated cells, **(b)** FFs-treated cells, and **(c)** RFs-treated cells. Cells were stained with BioTracker™ 555 Orange Cytoplasmic Membrane Dye (red) to assess plasma membrane morphology and integrity, and counterstained with DAPI (blue) to visualize cell nuclei. Images were acquired using an Invitrogen™ EVOS™ M7000 Imaging System at 20× magnification. Representative images demonstrate concentration-dependent alterations in plasma membrane morphology and nuclear staining following treatment with HEWL aggregates compared with untreated control cells. Scale bars represent 150 µm in all panels.

Cells treated with FFs exhibited mild concentration-dependent morphological alterations, including a reduction in the number of membrane-labeled cells and subtle changes in cell morphology, particularly at higher concentrations (Figure 6b). These observations are consistent with the modest decrease in metabolic activity detected by the XTT assay. In contrast, RF-treated cells exhibited pronounced morphological changes characterized by a marked reduction in BioTracker™ 555 fluorescence which indicates loss of normal cellular morphology compared with control and FF-treated cells (Figure 6c). These effects became progressively more evident with increasing RF concentration and are indicative of substantial membrane damage and cell loss. The fluorescence imaging findings closely corroborate the XTT viability results, demonstrating that rigid HEWL fibrils possess markedly higher neurotoxicity than flexible fibrils or native HEWL.

## 4 Discussion

The emergence of AMR has intensified the search for alternative antimicrobial materials that operate through mechanisms distinct from conventional antibiotics. In this study, we demonstrate that structural transformation of HEWL into amyloid fibrillar polymorphs significantly enhances its antibacterial activity against the Gram-negative bacteria *Escherichia coli* and *Salmonella* Typhimurium. Structural analyses confirmed the successful formation of two distinct fibrillar polymorphs, namely FFs and RFs, which differed in their secondary structure organization and morphology. CD spectroscopy, ThT fluorescence, and TEM collectively revealed that RFs possessed a more ordered β-sheet-rich architecture, whereas FFs exhibited a less compact and more flexible fibrillar network. These findings indicate that fibril formation conditions strongly influence the structural properties of HEWL amyloids and ultimately their biological activity. Native HEWL is a well-established antimicrobial protein whose antibacterial activity primarily arises from enzymatic hydrolysis of peptidoglycan within bacterial cell walls. However, this mechanism is generally more effective against Gram-positive bacteria, as the outer membrane of Gram-negative organisms restricts access to the underlying peptidoglycan layer [6]. Consistent with this limitation, native HEWL displayed only moderate antibacterial activity against both *E. coli* and *S.* Typhimurium. Remarkably, conversion into amyloid fibrillar assemblies substantially improved antibacterial efficacy, indicating that fibrillation introduces additional mechanisms of bacterial inhibition beyond the classical enzymatic function of lysozyme.

An equally important consideration for the development of antimicrobial biomaterials is their compatibility with mammalian cells. In the present study, native HEWL exhibited minimal cytotoxicity toward SH-SY5Y cells, whereas FFs caused only a modest reduction in cell viability and minor morphological changes. In contrast, RFs significantly reduced cell viability and induced pronounced membrane damage, indicating that fibril polymorphism strongly influences mammalian cell responses. The higher cytotoxicity of RFs is likely due to their more compact, highly ordered β-sheet-rich structure, which enhances membrane binding and disruption [15] [16]. In contrast, the more flexible and less densely packed architecture of FFs may reduce nonspecific interactions with mammalian membranes while still permitting effective interactions with negatively charged bacterial membranes [17]. Since bacterial membranes are enriched in anionic phospholipids, whereas mammalian membranes are predominantly zwitterionic and cholesterol-rich, FFs exhibit greater antibacterial selectivity with lower mammalian cytotoxicity [18].

The enhanced antibacterial activity of HEWL fibrils is likely associated with their highly ordered supramolecular architecture. Amyloid fibrils present repetitive arrays of charged and hydrophobic residues capable of interacting with bacterial membranes [19]. Electrostatic attraction between positively charged lysozyme residues and negatively charged bacterial surfaces may facilitate membrane association, while exposed hydrophobic domains can promote membrane destabilization and increased permeability. Such interactions resemble the membrane-targeting mechanisms reported for several antimicrobial peptides and functional amyloids [9]. Therefore, the antibacterial activity observed for the fibrillar polymorphs likely results from a combination of membrane interactions and residual lysozyme bioactivity. Among the fibrillar assemblies, FFs consistently exhibited stronger antibacterial activity than RFs. This observation highlights the importance of fibril morphology in determining biological function. In addition, FFs are formed under conditions that may preserve portions of the native protein structure, potentially retaining functional regions that contribute to antimicrobial activity. In contrast, the densely packed β-sheet architecture of RFs may reduce accessibility of surface residues, thereby limiting interactions with bacterial cells despite their higher structural order.

The observed morphology-dependent antibacterial activity supports the emerging concept that amyloid assemblies can serve as functional biomaterials rather than merely pathological protein aggregates. Recent studies have shown that flexible lysozyme amyloid nanoNETs exhibit potent antimicrobial activity. Similarly, the enhanced activity of FFs observed here suggests that fibril flexibility contributes to improved antibacterial efficacy [10]. Our findings extend this concept by showing that controlled modulation of HEWL fibril structure can significantly alter antibacterial performance. Importantly, the superior activity of FFs suggests that partially ordered and structurally dynamic amyloid assemblies may provide an optimal balance between stability and biological functionality. Overall, this study demonstrates that amyloid polymorphism is a key determinant of the antibacterial properties of HEWL assemblies. Structural tuning of lysozyme into flexible and rigid fibrillar forms generates materials with markedly enhanced activity against Gram-negative bacteria, with FFs exhibiting the greatest efficacy. These findings highlight the potential of engineered protein amyloids as bio-inspired antimicrobial materials and provide a framework for designing functional fibrillar biomaterials with tunable biological activities. Further studies are required to elucidate the molecular mechanisms responsible for the enhanced antibacterial activity of HEWL fibrils. Investigations of membrane permeability, fibril–membrane interactions, and bacterial ultrastructural changes following treatment will help distinguish the relative contributions of membrane disruption and residual enzymatic activity. In addition, evaluating these fibrillar assemblies in biologically relevant environments will be essential for assessing their potential applications as antimicrobial coatings, wound-healing materials, and therapeutic antimicrobial platforms.

## 5. Acknowledgements

We acknowledge the Biological Science Imaging Resource (BSIR) and the Institute of Molecular Biophysics (IMB) for access to core facilities. We thank Peter Randolph (IMB) and Anthony Warrington (BSIR) for their technical support.

## 6. Author Contributions

Sanjay Metkar: Investigation, Methodology, Data curation, Formal analysis, Writing – original draft and review & editing. Annie Scutte: Investigation, Data curation, Writing – review & editing. Jamel Ali: Investigation, Writing – review & editing. Ayyalusamy Ramamoorthy: Conceptualization, Supervision, Funding acquisition, Project administration, Writing – review & editing.

