## Supporting Information for "Structural Tuning of HEWL Amyloid Polymorphs Enhances Antibacterial Activity Against Gram-Positive and Gram-Negative Pathogens"

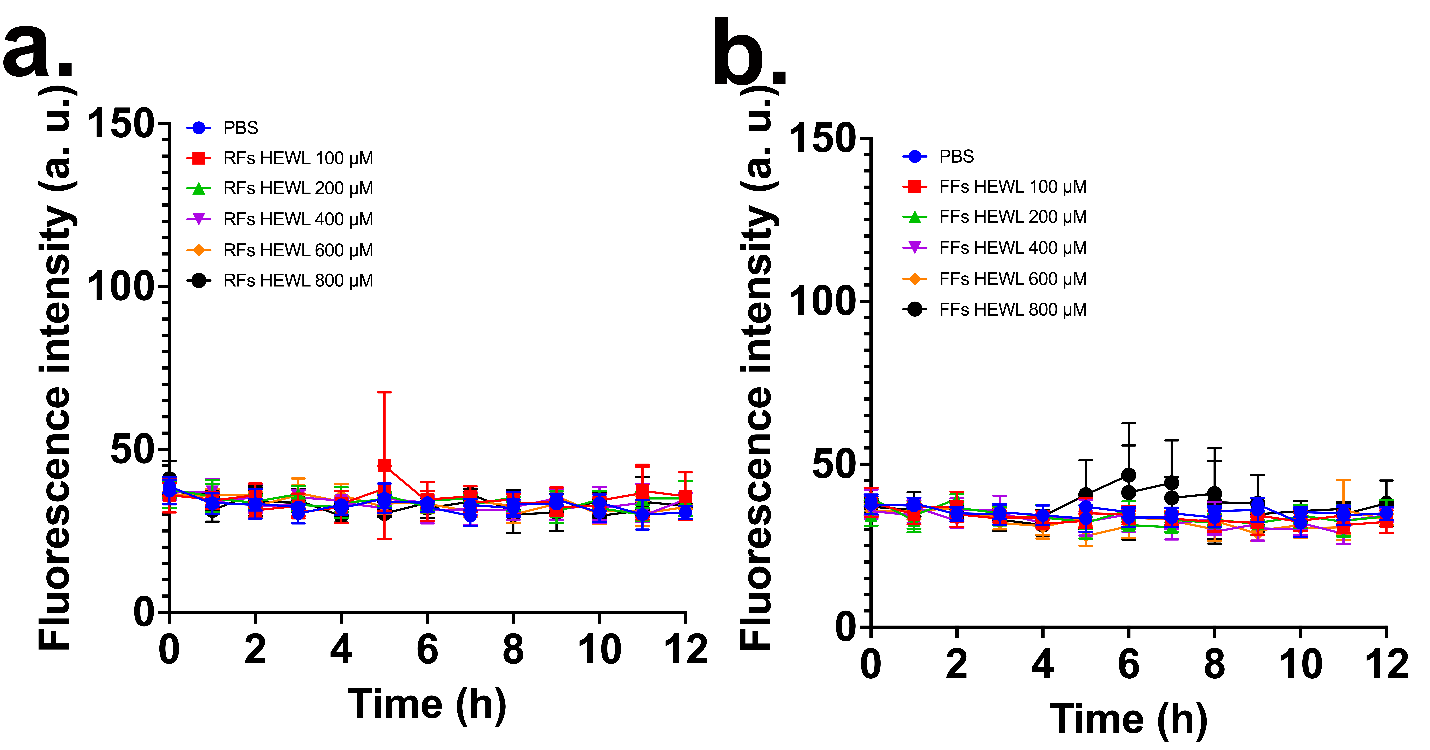

**Figure S1. Assessment of potential interference of HEWL amyloid fibrils with the Alamar Blue assay.** Fluorescence intensity of Alamar Blue in the presence of (a) RFs and (b) FFs of HEWL at concentrations ranging from 100–800 μM in the absence of bacteria. PBS served as the control. Fluorescence signals remained comparable to the PBS control throughout the 12 h incubation period, indicating that neither RFs nor FFs directly reduced resazurin or generated significant background fluorescence. These results confirm that the antibacterial effects observed in Figures 2–4 arise from bacterial growth inhibition rather than interference of the fibrillar materials with the Alamar Blue assay. Data are presented as mean ± SD (n = 3).
